# Correlation, Regression, and Growth Curve Analyses of BodyWeight and Body Measurements in Female Jiangyue Donkeys

**DOI:** 10.64898/2026.04.28.721092

**Authors:** Wei Ren, Lei Zhao, Huaixing Yin, Qiong Wang, Lingling Liu

## Abstract

**Objective:** This study aimed to systematically analyze the correlation between body weight and body measurement traits and the growth pattern of female Jiangyue donkeys.

**Methods:** A total of 484 female Jiangyue donkeys were selected to determine their body weight (Y) and 11 body measurement traits, including withers height (X_1_), body length (X_2_), chest circumference (X_3_), cannon circumference (X_4_), head length (X_5_), neck length (X_6_), chest width (X_7_), chest depth (X_8_), rump height (X_9_), rump length (X_10_) and rump width (X_11_). SPSS 27.0 statistical software was used to conduct descriptive statistics, correlation and regression analysis on body weight and body measurement traits, and the optimal regression equation between them was established by the stepwise method and validated. Furthermore, four growth curve models (Logistic, Gompertz, Brody and Von Bertalanffy) were used to fit the body weight of 241 female Jiangyue donkeys at different months of age.

**Results:** The body weight of female Jiangyue donkeys was significantly positively correlated with all body measurement traits (p<0.01), with the highest correlation coefficient observed for chest circumference. The optimal regression equations for body weight established by stepwise regression yielded R² values of 0.918 and 0.844 for growing and adult donkeys, respectively (p<0.01). Among the four growth curve models, the Von Bertalanffy model exhibited the best fitting effect (R²=0.99992), with an estimated asymptotic body weight of 184.41 kg, which was close to the measured values of growing female donkeys.

**Conclusion:** The regression models for body weight estimation and the Von Bertalanffy growth curve model for growth pattern evaluation established in this study can serve as effective tools and production management of female Jiangyue donkeys.

## INTRODUCTION

China has a history of donkey raising for more than 4,000 years and owns 24 local donkey breeds. Up to now, the donkey industry remains an indispensable and important part of China’s characteristic animal husbandry [1,2,3]. With the development of the times and advances in science and technology, the draught value of donkeys as livestock has been declining, leading to an overall downward trend in China’s donkey stock [4]. Data from the National Bureau of Statistics of China in 2025 showed that China’s donkey stock dropped from 7.7716 million heads in 2005 to 1.2844 million heads in 2024, with a decrease of about 83.47% in two decades [5]. At present, four donkey breeds are at risk of endangerment, reflecting the systematic challenges faced by the protection of donkey germplasm resources and the industrial development of the donkey industry in China [6]. Nowadays, the functions of donkeys have gradually shifted to the production of donkey milk, donkey hide, donkey meat and recreational riding, and the consumer demand for donkey products has been continuously growing [7].

Since the 1970s, Guanzhong donkeys, a large donkey breed from Shaanxi Province, have been gradually introduced to Xinjiang for crossbreeding with local small Xinjiang donkeys, thus breeding Jiangyue donkey, a characteristic and excellent breed with the advantages of strong stress resistance, roughage tolerance and high production performance [8]. Jiangyue donkeys are mainly distributed in Kashgar, Hotan, and other parts of Xinjiang. They have not yet been officially recognized as a standardized breed and are therefore better regarded as a cultivated commercial population [9]. Against this background, the present study analyzed the relationships between body weight and body measurements in female Jiangyue donkeys at different growth stages and compared several nonlinear models to describe age-related body-weight changes.

## MATERIALS AND METHODS

### Experimental Animals

The data were provided by Xinjiang Jinhuyang Animal Husbandry Co., Ltd. and derived from two donkey populations: a total of 484 female donkeys were used for the correlation and regression analysis of body measurements and body weight, including 296 growing donkeys (7 months to 3 years old) and 188 adult donkeys (4 to 13 years old). Another 241 female donkeys at different months of age (1 to 14 months) were selected for growth curve fitting. Among these, individuals aged 1 to 6 months (n=120) were not included in the regression analysis, while those aged 7 to 14 months (n=121) were part of the growing donkey group (7 months to 3 years old) used for regression. Thus, the growth curve and regression datasets partially overlap for the 7 to 14 month age range. All experimental donkeys were raised under the same environmental and facility conditions and received standardized feeding and management procedures of the enterprise to ensure consistent management conditions.

### Measurement Methods and Indicators

Body weight (Y) before feeding was measured using a platform scale with an accuracy of 0.5 kg, and body measurements were determined with a soft tape measure and a measuring stick (both with an accuracy of 0.1 cm). The body measurement indicators included withers height (X_1_), body length (X_2_), chest circumference (X_3_), cannon circumference (X_4_), head length (X_5_), neck length (X_6_), chest width (X_7_), chest depth (X_8_), rump height (X_9_), rump length (X_10_) and rump width (X_11_). The determination of body measurement indicators followed the measurement standards specified in Technical Specification for Donkey Production Performance Determination [10], and the specific measurement methods are shown in Table 1. To reduce human error and ensure measurement accuracy, all the above indicators were measured by the same person.

**Table 1.**
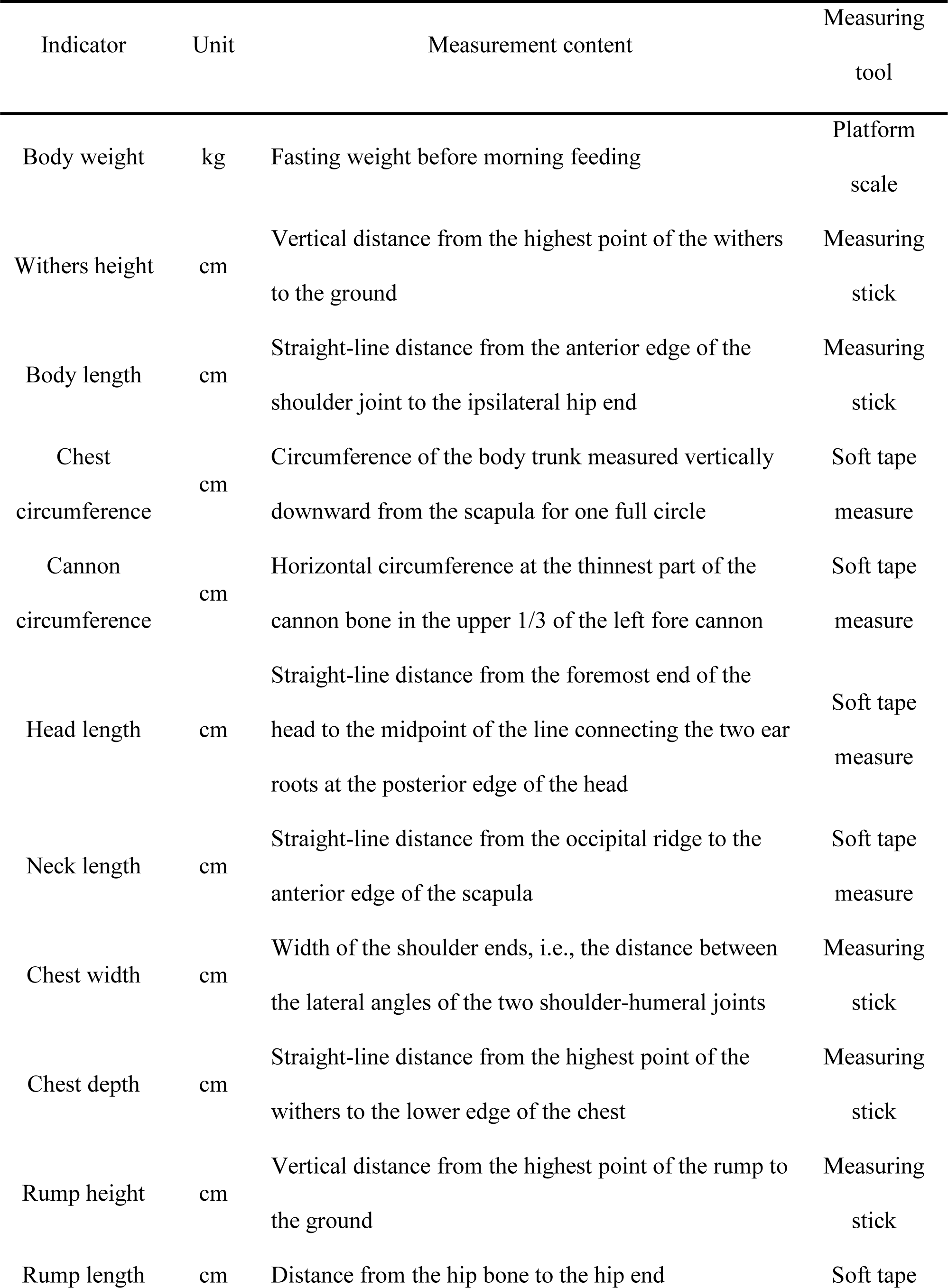
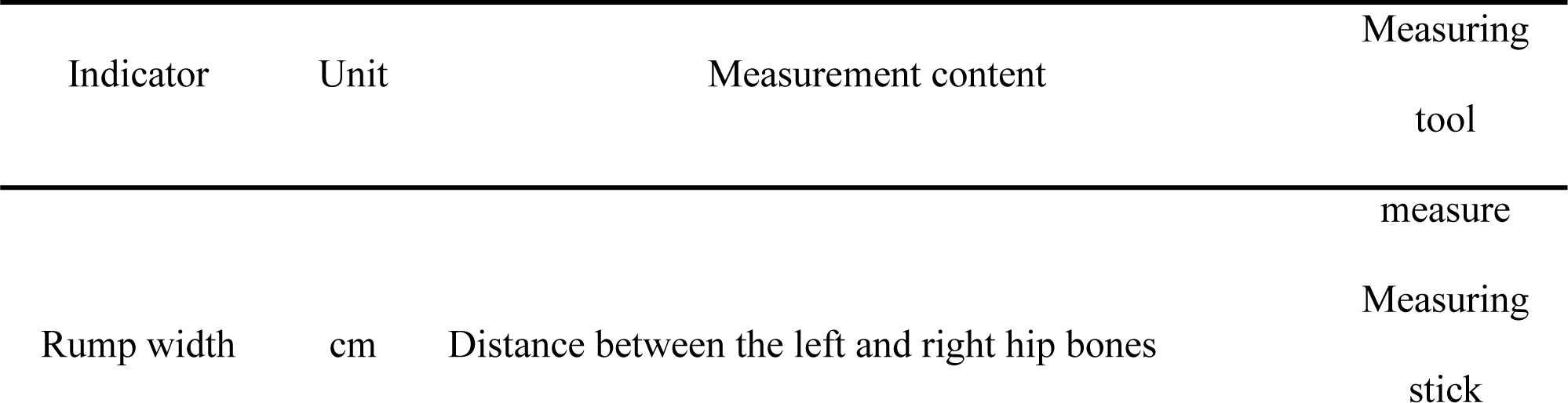
Measurement methods of body weight and body measurement indicators.

### Statistical Analysis

After sorting the measured data in Excel, SPSS 27.0 statistical software was used to conduct descriptive statistics and correlation analysis on 484 female Jiangyue donkeys at different growth stages, and the stepwise method was adopted to screen out the optimal linear regression models between body weight and various body measurement traits of female Jiangyue donkeys at different growth stages. Origin 2024 software was used to fit the body weight of 241 female Jiangyue donkeys at different months of age with four curve models (Logistic, Gompertz, Brody and Von Bertalanffy), and the equations of each model are shown in Table 2. The fitting curves and actual growth curves were also plotted.

**Table 2.**
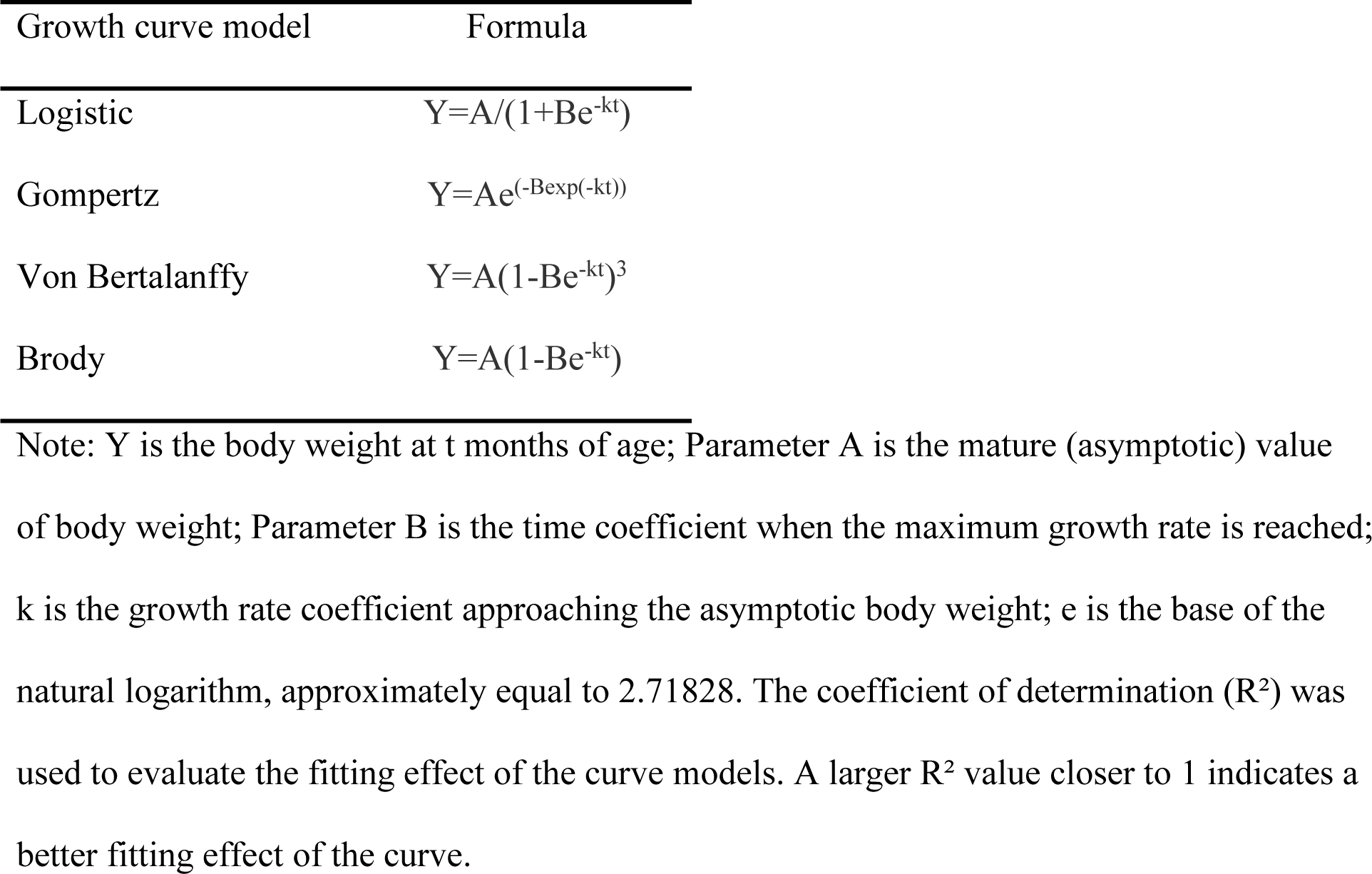
Different growth curve models Growth curve model Formula Logistic Y=A/(1+Be^-kt^)

## RESULTS

### Descriptive Statistical Analysis of Body Weight and Body Measurements

Descriptive statistical analysis was conducted on 296 growing female Jiangyue donkeys (hereinafter referred to as growing donkeys) and 188 adult female Jiangyue donkeys (hereinafter referred to as adult donkeys), and the results are shown in Table 3 and Table 4. The higher the coefficient of variation (CV), the greater the degree of dispersion of the data, indicating more pronounced individual differences.The CV of body weight and body measurement traits of growing donkeys ranged from 4.80% to 19.04%, with the CV of body weight being the highest (19.04%). For the other 11 body measurement traits, the CVs in descending order were rump width, neck length, rump length, chest width, body length, cannon circumference, chest circumference, chest depth, head length, withers height and rump height. The CV of body weight and body measurement traits of adult donkeys ranged from 5.62% to 16.00%, with the CV of body weight still being the highest (16.00%). The CVs of the other 11 body measurement traits in descending order were chest width, neck length, cannon circumference, chest depth, rump width, rump length, body length, head length, withers height, rump height and chest circumference.

**Table 3.** Phenotypic statistics of body weight and body measurements of growing female Jiangyue donkeys.

| Trait | Mean | Standard<br>deviation | Coefficient<br>of variation | Minimum | Maximum |
| --- | --- | --- | --- | --- | --- |
| Body weight (kg) | 173.86 | 33.103 | 19.04% | 89 | 273 |
| Withers height (cm) | 121.70 | 5.948 | 4.89% | 105 | 139 |
| Body length (cm) | 122.24 | 8.135 | 6.65% | 103 | 146 |
| Chest circumference<br>(cm) | 124.74 | 8.211 | 6.58% | 100 | 147 |
| Cannon circumference<br>(cm) | 14.14 | 0.936 | 6.62% | 12 | 18 |
| Head length (cm) | 49.20 | 3.007 | 6.11% | 42 | 58 |
| Neck length (cm) | 48.76 | 3.936 | 8.07% | 36 | 61 |
| Chest width (cm) | 28.62 | 1.979 | 6.91% | 23 | 35 |
| Chest depth (cm) | 52.91 | 3.250 | 6.14% | 44 | 62 |
| Rump height (cm) | 125.15 | 6.002 | 4.80% | 107 | 143 |
| Rump length (cm) | 37.65 | 2.689 | 7.14% | 30 | 45 |
| Rump width (cm) | 36.89 | 3.401 | 9.22% | 23 | 47 |

**Table 4.** Phenotypic statistics of body weight and body measurements of adult female Jiangyue donkeys.

| Trait | Mean | Standard<br>deviation | Coefficient<br>of variation | Minimum | Maximum |
| --- | --- | --- | --- | --- | --- |
| Body weight (kg) | 228.02 | 36.479 | 16.00% | 145 | 321 |
| Withers height (cm) | 125.88 | 7.223 | 5.74% | 109 | 146 |
| Body length (cm) | 130.26 | 8.301 | 6.37% | 100 | 164 |
| Chest circumference<br>(cm) | 136.81 | 7.693 | 5.62% | 117 | 155 |
| Cannon circumference<br>(cm) | 14.41 | 1.121 | 7.78% | 12 | 18 |
| Head length (cm) | 52.38 | 3.079 | 5.88% | 42 | 61 |
| Neck length (cm) | 53.56 | 4.761 | 8.89% | 42 | 66 |
| Chest width (cm) | 29.81 | 2.731 | 9.16% | 22 | 38 |
| Chest depth (cm) | 57.20 | 4.210 | 7.36% | 48 | 70 |
| Rump height (cm) | 129.13 | 7.320 | 5.67% | 109 | 148 |
| Rump length (cm) | 39.73 | 2.666 | 6.71% | 30 | 47 |
| Rump width (cm) | 39.70 | 2.767 | 6.97% | 30 | 48 |

### Correlation Analysis of Body Weight and Body Measurements

To explore the relationship between body weight and body measurements of female Jiangyue donkeys at different growth stages, correlation analysis was conducted on the 12 traits. The results, as shown in Table 5 and Table 6, indicated that the body weight of growing and adult donkeys was significantly positively correlated with all body measurement traits (p<0.01), with the correlation coefficient between body weight and chest circumference being the highest among all traits, at 0.929 and 0.904, respectively. For growing donkeys, the correlation coefficients of other body measurement traits with body weight in descending order were rump length (0.851), rump width (0.838), rump height (0.787), body length (0.776), withers height (0.762), chest width (0.719), head length (0.652), cannon circumference (0.627), chest depth (0.540) and neck length (0.517). Among the 11 body measurement indicators of growing donkeys, the correlation coefficient between withers height and rump height was 0.924, indicating a strong positive correlation between them; in contrast, the correlation coefficient between neck length and cannon circumference was only 0.264, showing a weak correlation. For adult donkeys, the correlation coefficients of other body measurement traits with body weight in descending order were rump height (0.767), withers height (0.748), body length (0.736), rump width (0.732), rump length (0.695), cannon circumference (0.625), chest width (0.589), chest depth (0.554), neck length (0.502) and head length (0.443). Among the 11 body measurement indicators of adult donkeys, the correlation coefficient between withers height and rump height was 0.950, indicating a strong positive correlation, which was consistent with the result of growing donkeys; on the contrary, the correlation coefficient between head length and neck length was 0.240, indicating a weak correlation. All correlation coefficients presented above are Pearson correlation coefficients.

**Table 5.**
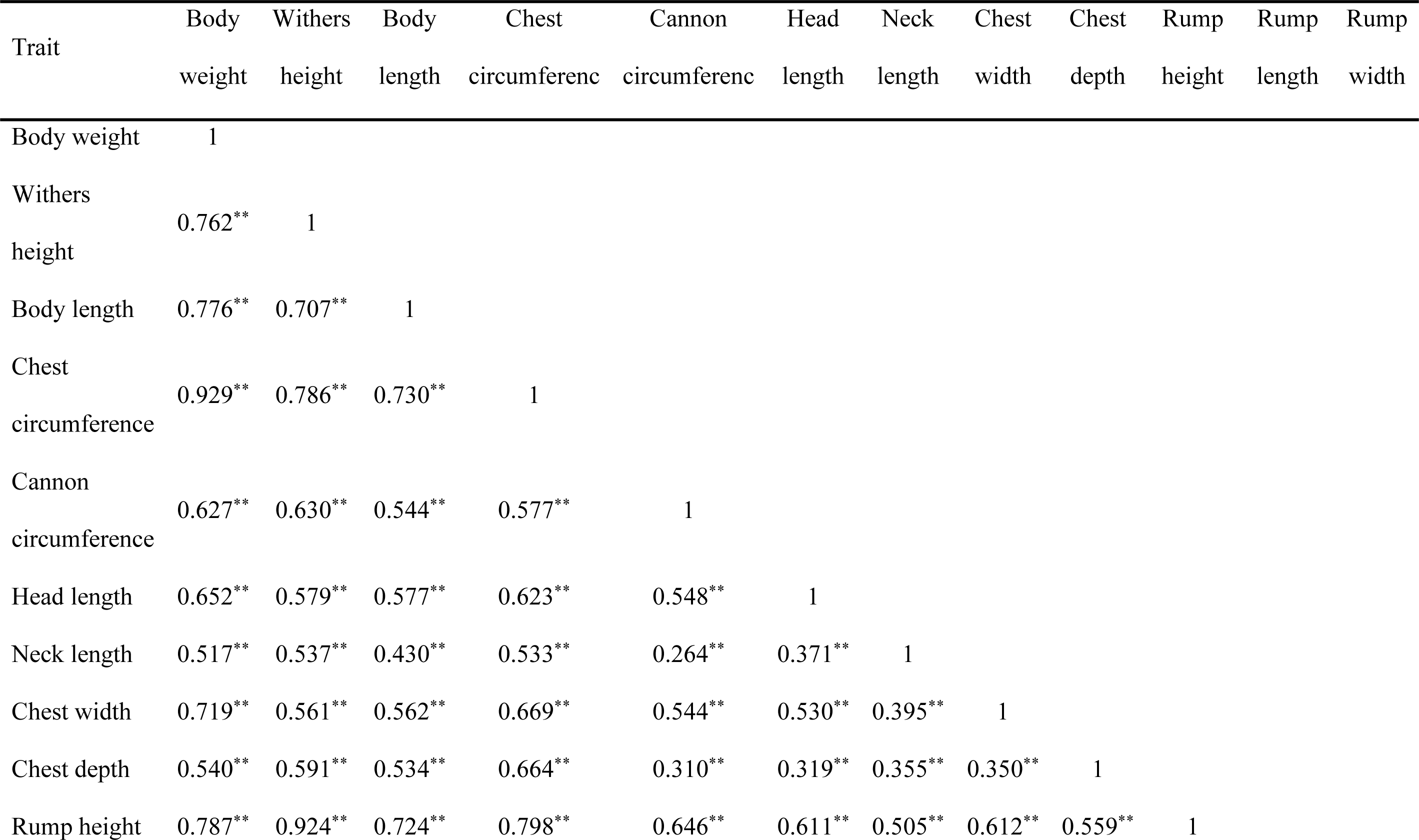
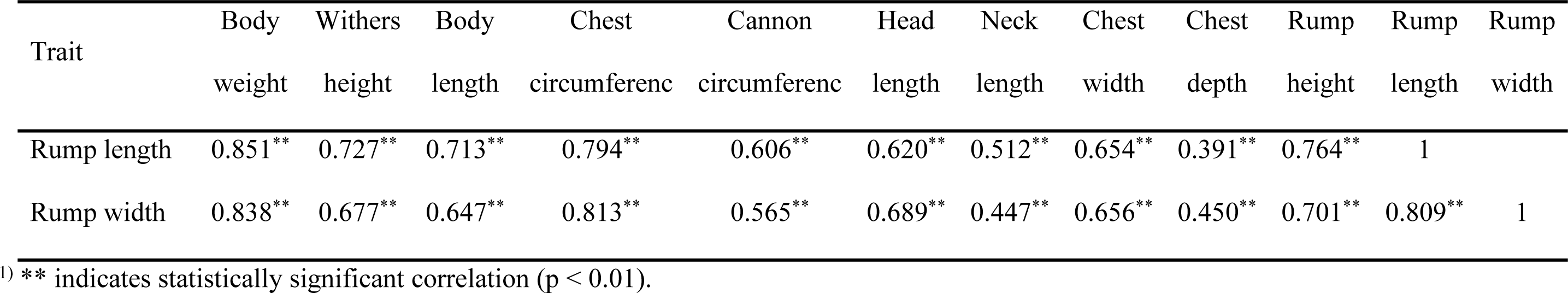
Correlation coefficients between body weight and body measurement indicators of growing female Jiangyue donkeys.

**Table 6.**
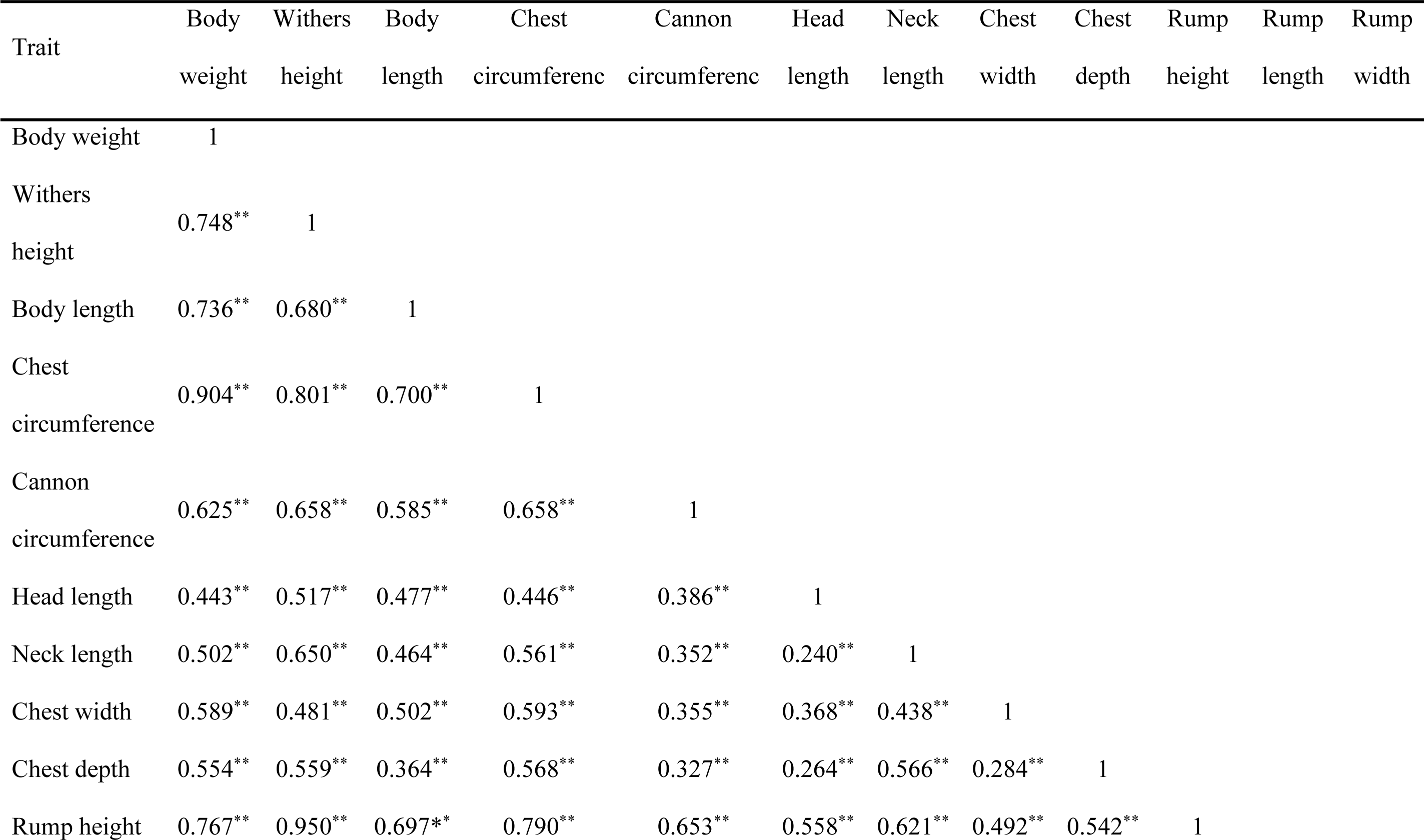
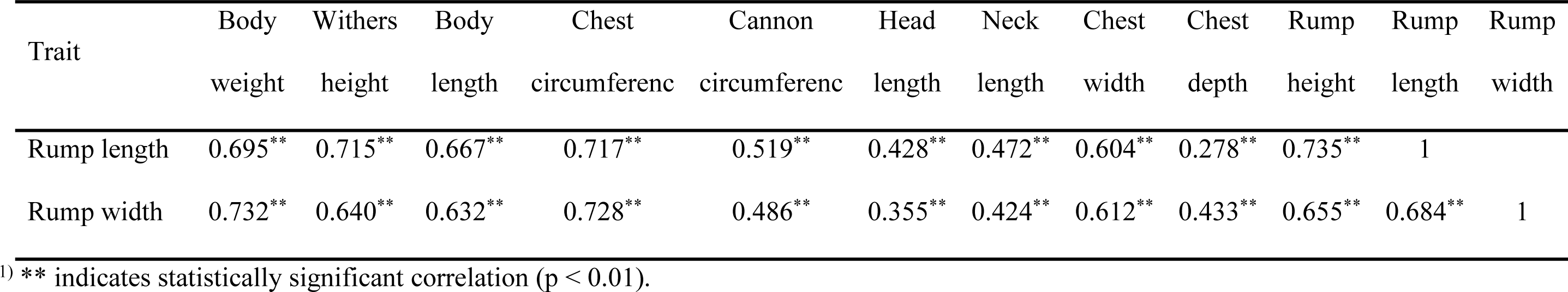
Correlation coefficients between body weight and body measurement indicators of adult female Jiangyue donkeys.

| Trait | Body weight | Withers height | Body length | Chest circumference | Cannon circumference | Head length | Neck length | Chest width | Chest depth | Rump height | Rump length | Rump width |
| --- | --- | --- | --- | --- | --- | --- | --- | --- | --- | --- | --- | --- |
| Body weight | 1 |  |  |  |  |  |  |  |  |  |  |  |
| Withers height | 0.748** | 1 |  |  |  |  |  |  |  |  |  |  |
| Body length | 0.736** | 0.680** | 1 |  |  |  |  |  |  |  |  |  |
| Chest circumference | 0.904** | 0.801** | 0.700** | 1 |  |  |  |  |  |  |  |  |
| Cannon circumference | 0.625** | 0.658** | 0.585** | 0.658** | 1 |  |  |  |  |  |  |  |
| Head length | 0.443** | 0.517** | 0.477** | 0.446** | 0.386** | 1 |  |  |  |  |  |  |
| Neck length | 0.502** | 0.650** | 0.464** | 0.561** | 0.352** | 0.240** | 1 |  |  |  |  |  |
| Chest width | 0.589** | 0.481** | 0.502** | 0.593** | 0.355** | 0.368** | 0.438** | 1 |  |  |  |  |
| Chest depth | 0.554** | 0.559** | 0.364** | 0.568** | 0.327** | 0.264** | 0.566** | 0.284** | 1 |  |  |  |
| Rump height | 0.767** | 0.950** | 0.697** | 0.790** | 0.653** | 0.558** | 0.621** | 0.492** | 0.542** | 1 |  |  |

| Trait | Body<br>weight | Withers<br>height | Body<br>length | Chest<br>circumferenc | Cannon<br>circumferenc | Head<br>length | Neck<br>length | Chest<br>width | Chest<br>depth | Rump<br>height | Rump<br>length | Rump<br>width |
| --- | --- | --- | --- | --- | --- | --- | --- | --- | --- | --- | --- | --- |
| Rump length | 0.695** | 0.715** | 0.667** | 0.717** | 0.519** | 0.428** | 0.472** | 0.604** | 0.278** | 0.735** | 1 |  |
| Rump width | 0.732** | 0.640** | 0.632** | 0.728** | 0.486** | 0.355** | 0.424** | 0.612** | 0.433** | 0.655** | 0.684** | 1 |

<sup>1)</sup> \*\* indicates statistically significant correlation ( $p < 0.01$ ).

### Stepwise Regression Analysis of Body Weight and Body Measurements

SPSS 27.0 statistical software was used to establish a multiple linear regression equation with body weight as the dependent variable and 11 body measurement traits as independent variables, and the corresponding models were obtained by the stepwise analysis method, as shown in Table 7. The main body measurement indicators included in the regression equation for growing donkeys were body length (X_2_), chest circumference (X_3_), chest width (X_7_), chest depth (X_8_), rump length (X_10_) and rump width (X_11_), while those for adult donkeys were body length (X_2_), chest circumference (X_3_) and rump width (X_11_). Further analysis of the equation coefficients of the two regression models was conducted, and the results are shown in Table 8. Combined with Table 7 and Table 8, the optimal regression equation for body weight and body measurement traits of growing donkeys was Y=0.614X_2_+2.501X_3_+1.472X_7_-0.863X_8_+1.807X_10_+0.953X_11_-312.925 (Equation 1), and that for adult donkeys was Y=0.774X_2_+3.320X_3_+1.461X_11_-385.023 (Equation 2).

**Table 7.**
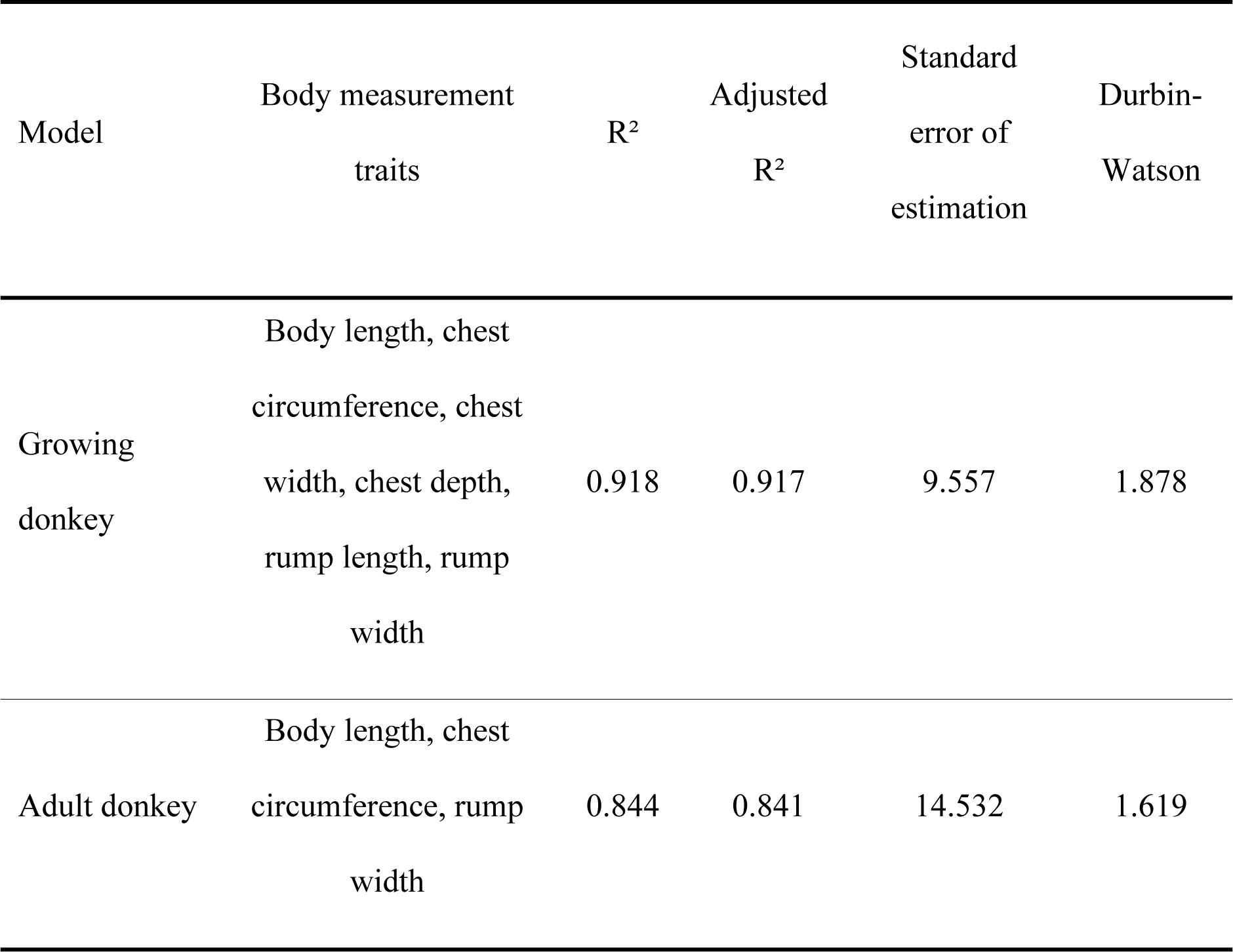
Regression models of body weight on body measurement traits of female Jiangyue.

| Model | Body measurement traits | R<sup>2</sup> | Adjusted R<sup>2</sup> | Standard error of estimation | Durbin-Watson |
| --- | --- | --- | --- | --- | --- |
| Growing donkey | Body length, chest |  |  |  |  |
|  | circumference, chest |  |  |  |  |
|  | width, chest depth, | 0.918 | 0.917 | 9.557 | 1.878 |
|  | rump length, rump width |  |  |  |  |
| Adult donkey | Body length, chest |  |  |  |  |
|  | circumference, rump | 0.844 | 0.841 | 14.532 | 1.619 |
|  | width |  |  |  |  |

**Table 8.**
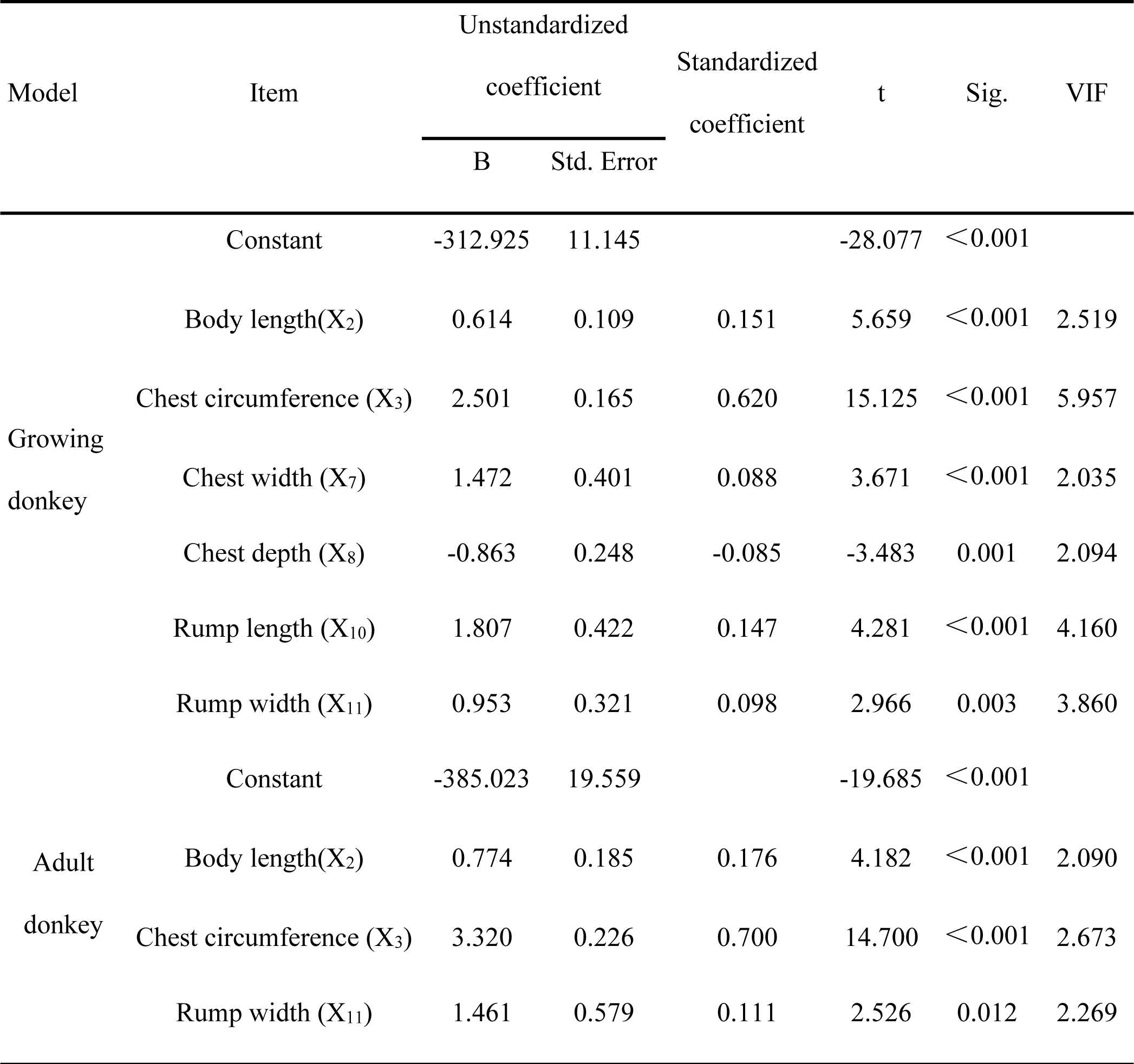
Regression coefficient analysis of regression equation models for female Jiangyue donkeys at different growth stages.

The optimal regression models explained 91.8% and 84.4% of the variance in body weight in growing and adult donkeys, respectively. Analysis of variance (ANOVA) was performed, and the results are shown in Table 9. The F-values of the two regression models were F_1_=541.738 and F_2_=331.459, respectively, both reaching a statistically significant level (p<0.01), which indicated that the selected body measurement traits jointly explained a substantial proportion of the variation in body weight.

**Table 9.** ANOVA of stepwise regression between body weight and body measurements of female Jiangyue donkeys at different growth stages.

| Model | Item | Sum of<br>squares | df | Mean<br>square | F | Sig. |
| --- | --- | --- | --- | --- | --- | --- |
| Growing<br>donkey | Regression | 296873.209 | 6 | 49478.868 | 541.738 | <0.001 |
|  | Residual | 26395.386 | 289 | 91.334 |  |  |
|  | Total | 323268.595 | 295 |  |  |  |
| Adult<br>donkey | Regression | 209991.258 | 3 | 69997.086 | 331.459 | <0.001 |
|  | Residual | 38856.884 | 184 | 211.179 |  |  |
|  | Total | 248848.142 | 187 |  |  |  |

### Residual Analysis and Model Diagnosis

To verify whether the established optimal regression models conformed to the basic assumptions of linear regression, SPSS 27.0 was used to plot the P-P plots of standardized residuals, as shown in Figure 1 and Figure 2. The scatter points were distributed along and near the diagonal line, indicating that the residuals of the two regression models appeared approximately normally distributed. Figure 3 and Figure 4 showed that the standardized residuals were evenly distributed on both sides of the standardized predicted values, suggesting that the residuals of both models exhibited no obvious heteroscedasticity. The Durbin-Watson test was used to detect the autocorrelation of the data, and its statistical value ranged from 0 to 4. A value less than 2 indicates positive autocorrelation, a value greater than 2 indicates negative autocorrelation, and a value close to 2 means no autocorrelation (i.e., the data are independent of each other) [11]. As shown in Table 7, the Durbin-Watson values of the growing donkey model and the adult donkey model were 1.878 and 1.619, respectively, both close to 2, indicating that there was no obvious autocorrelation in the residuals of the two regression equations, i.e., no serial correlation among the data. The variance inflation factors (VIFs) for all predictors were below 6, indicating no severe multicollinearity. All the above test results confirmed that the established optimal regression equations were statistically significant.

**Figure 1.**
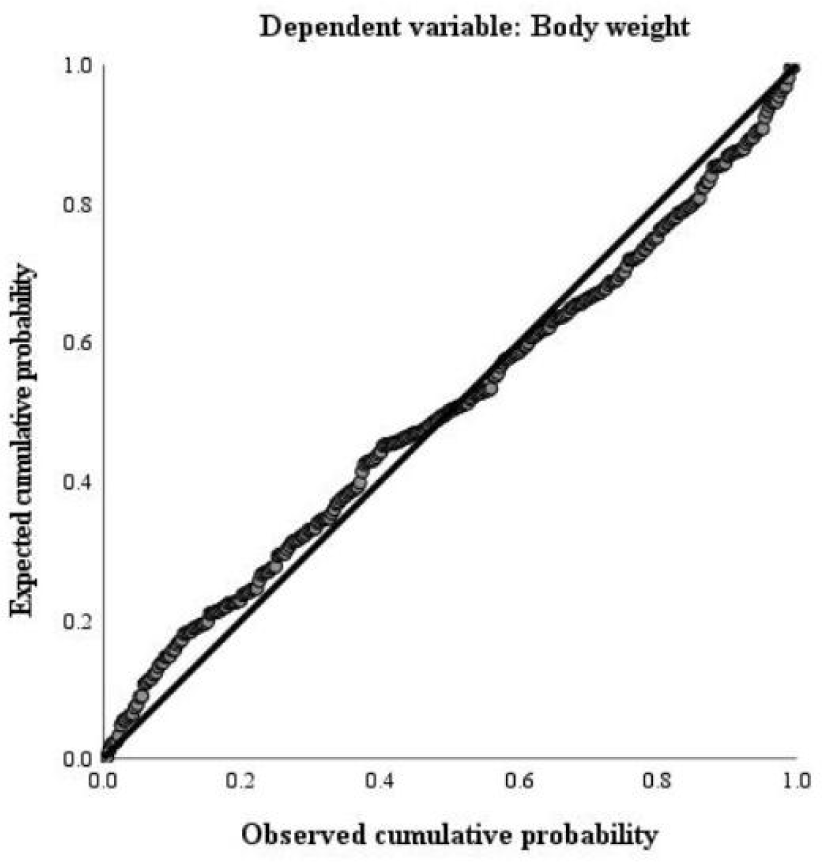
P-P plot of standardized residuals for body weight regression of growing female Jiangyue donkeys

**Figure 2.**
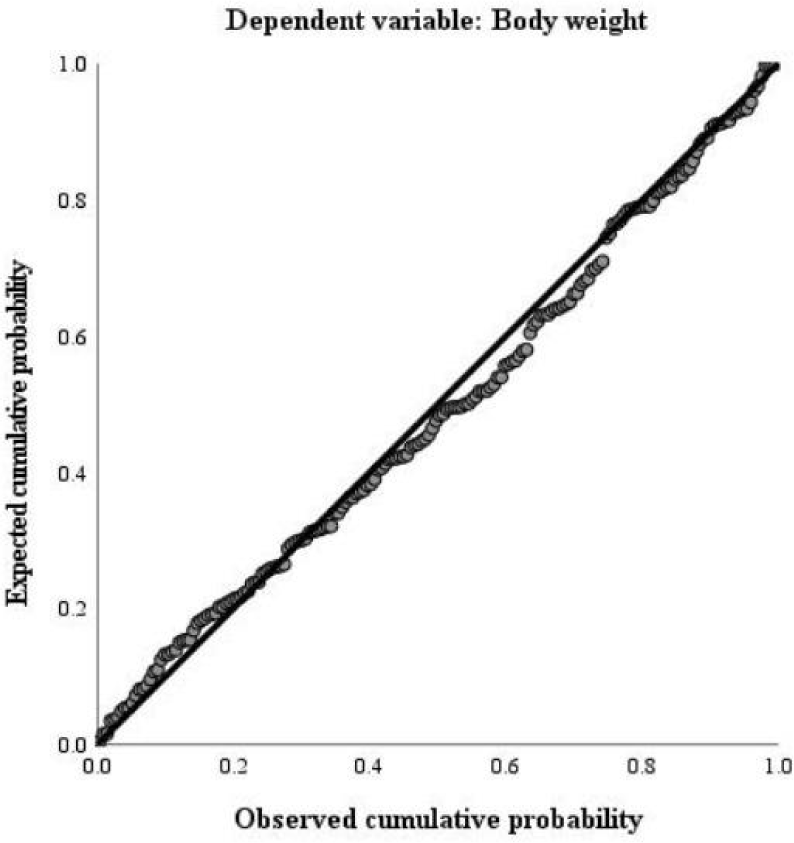
P-P plot of standardized residuals for body weight regression of adult female Jiangyue donkeys

**Figure 3.**
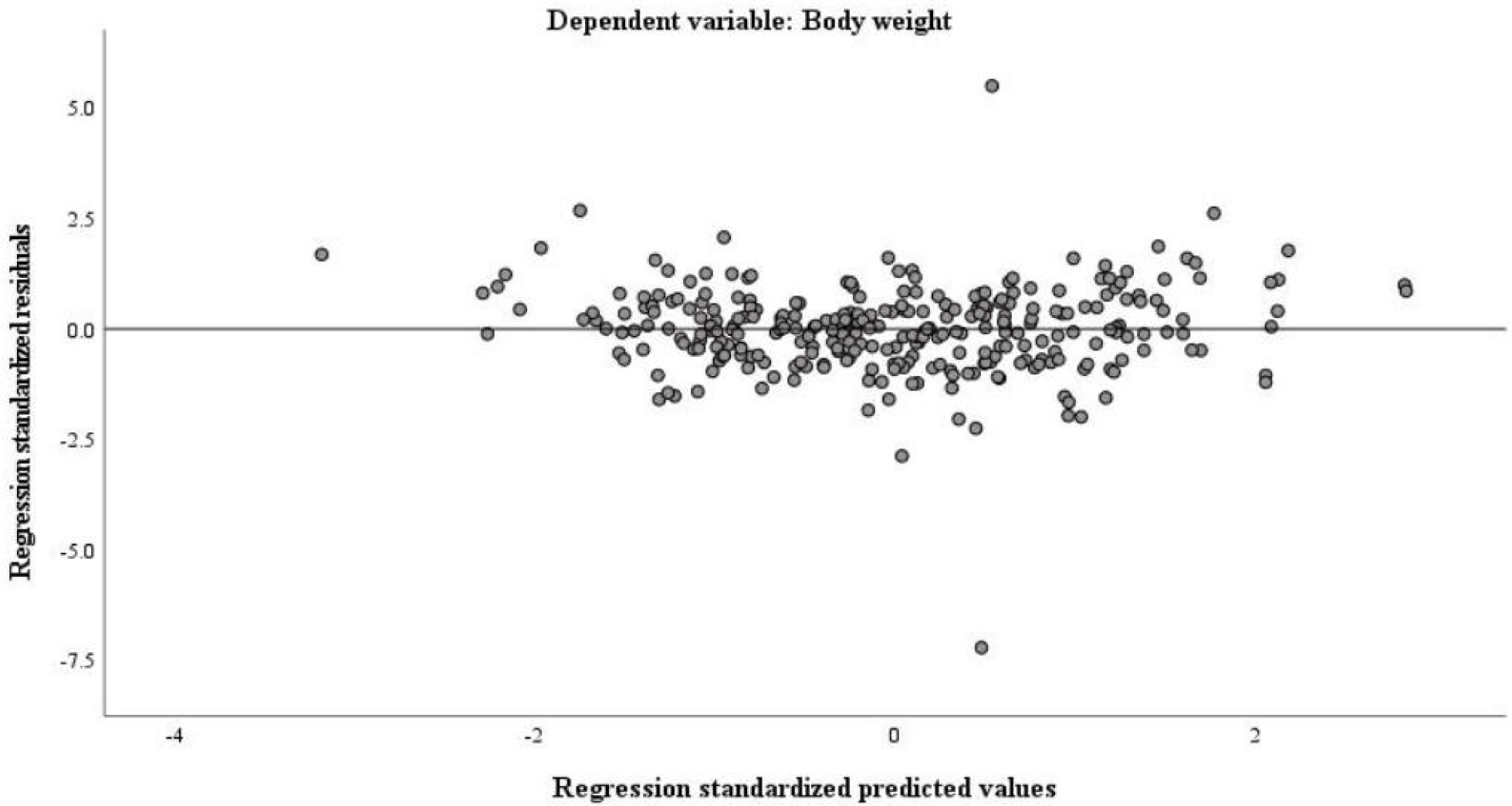
Scatter plot of standardized residuals for body weight regression of growing female Jiangyue donkeys

**Figure 4.**
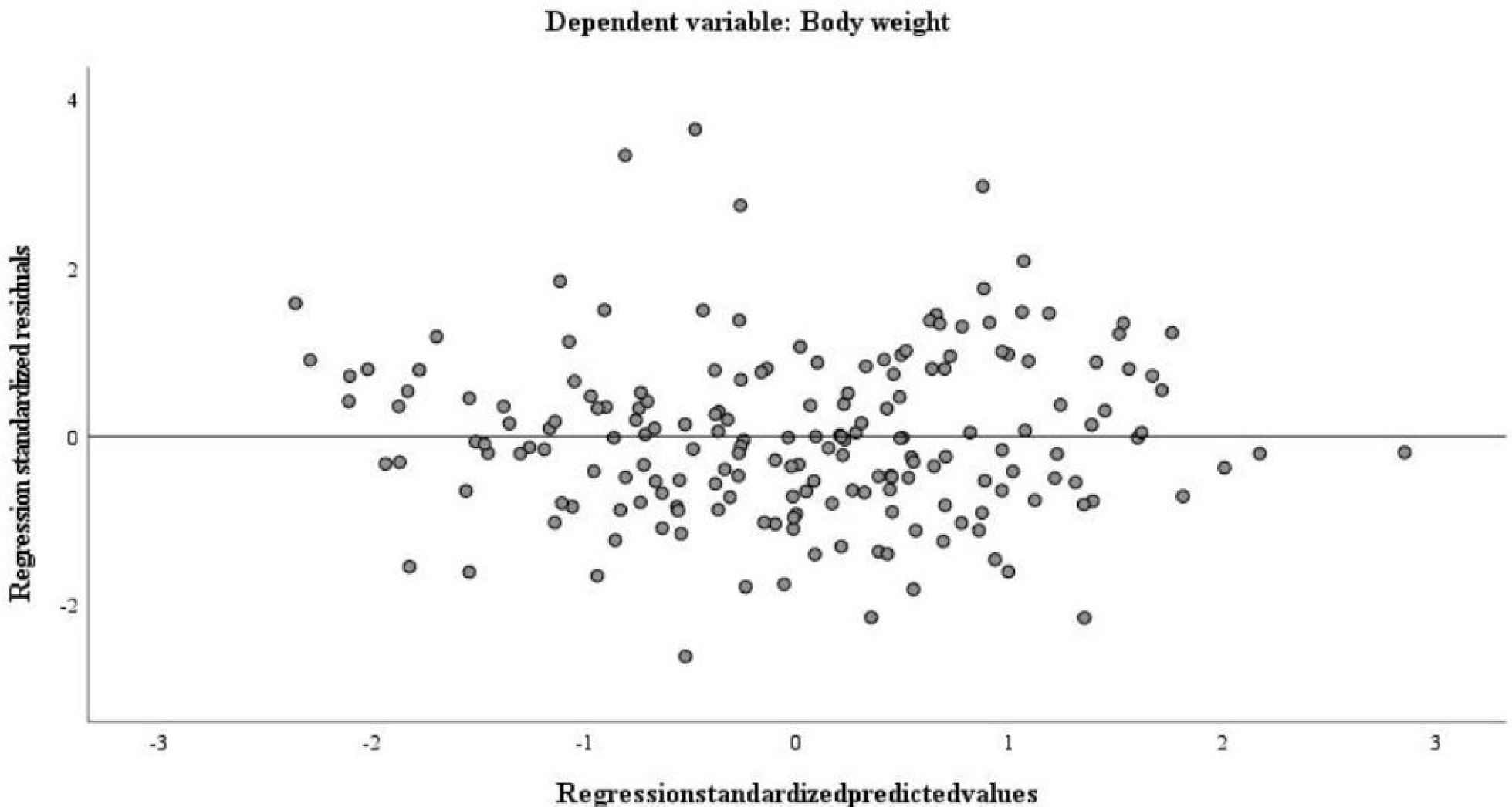
Scatter plot of standardized residuals for body weight regression of adult female Jiangyue donkeys

### Validation of Regression Models

To avoid overfitting, the growing donkey data were randomly split into a training set (n=236) and a test set (n=60) at a ratio of 8:2 using a random splitting method. Stepwise regression analysis was re-performed on the training set, and the selected variables were consistent with those of the full sample. The regression equation was: Y=0.581X_2_+2.546X_3_+1.480X_7_-0.710X_8_+1.876X_10_+0.768X_11_-317.939(R²=0.915,p<0.01), which was highly consistent with the result from the full sample (R²=0.918). The model developed from the training set was applied to the test set, and the prediction error metrics are shown in Table 10. For the test set, the R² was 0.933, with an RMSE of 8.32 kg, an MAE of 6.83 kg, and a MAPE of 4.28%. The R² of the test set was slightly higher than that of the training set, which may be attributed to fluctuations in the sample distribution of the test set following random splitting. Such minor fluctuations are normal when the sample size is limited. Using the same method, the adult donkey data were split into a training set (n=146) and a test set (n=42) at a ratio of 8:2. The stepwise regression model from the training set included the same variables as those in the full sample, and the equation was: Y=0.648X_2_+3.428X_3_+1.511X_11_-385.956(R²=0.843,p<0.01), which was highly consistent with the full sample result (R²=0.844). This model was then applied to the test set, yielding an R² of 0.847, an RMSE of 13.71 kg, an MAE of 11.88 kg, and a MAPE of 5.39%. The R² values of the test set and training set were similar, indicating that the model’s explanatory ability on unseen data was basically consistent with that on the modeling data, and no obvious overfitting occurred. Overall, all error metrics were low, demonstrating the model’s good generalization ability. The similar performance between the training and test sets suggests that the models were relatively stable and did not indicate obvious overfitting under the current data split. Thus, the full-sample estimated equation was used as the final model.

**Table 10.** Validation results of the body weight regression models for Jiangyue donkeys.

| Population | Sample size<br>(training/test) | R<sup>2</sup><br>(training) | R<sup>2</sup> (test) | RMSE<br>(kg) | MAE<br>(kg) | MAPE<br>(%) |
| --- | --- | --- | --- | --- | --- | --- |
| Growing<br>donkeys | 236 / 60 | 0.915 | 0.933 | 8.32 | 6.83 | 4.28 |
| Adult<br>donkeys | 146 / 42 | 0.843 | 0.847 | 13.71 | 11.88 | 5.39 |

### Growth Curve Fitting with Different Models

In this study, body weight traits of Jiangyue donkeys were determined at 1–14 months of age. Valid age points included in the growth curve fitting were 1, 2, 3, 4, 7, 8, 9, 10, 12 and 14 months, with the remaining age points excluded owing to inadequate valid sample sizes. The average body weight and standard deviation of 241 female Jiangyue donkeys at different months of age are shown in Table 11. Four curve models (Logistic, Gompertz, Brody and Von Bertalanffy) were used to fit their average body weight. Combined with Table 12 and Figure 5, the body weight of female Jiangyue donkeys increased rapidly before 4 months of age, and then the growth rate gradually slowed down. All four curve models exhibited good fits for body weight, with R² values all above 0.96418, and all models showed statistically significant fits (p<0.01), which captured the age-related trend in body weight reasonably well. Among the four models, the Von Bertalanffy model had the highest R² value (0.99992) for fitting the body weight of female Jiangyue donkeys, which was higher than those of the Logistic, Gompertz and Brody models. It could be seen that the Von Bertalanffy model had the best fitting effect on the body weight of female Jiangyue donkeys, with an asymptotic body weight of 184.41 kg. Within the observed age range, the Von Bertalanffy model provided the best fit among the four candidate models. However, the estimated asymptotic weight should be interpreted with caution because the data covered only the early growth stage.

**Figure 5.**
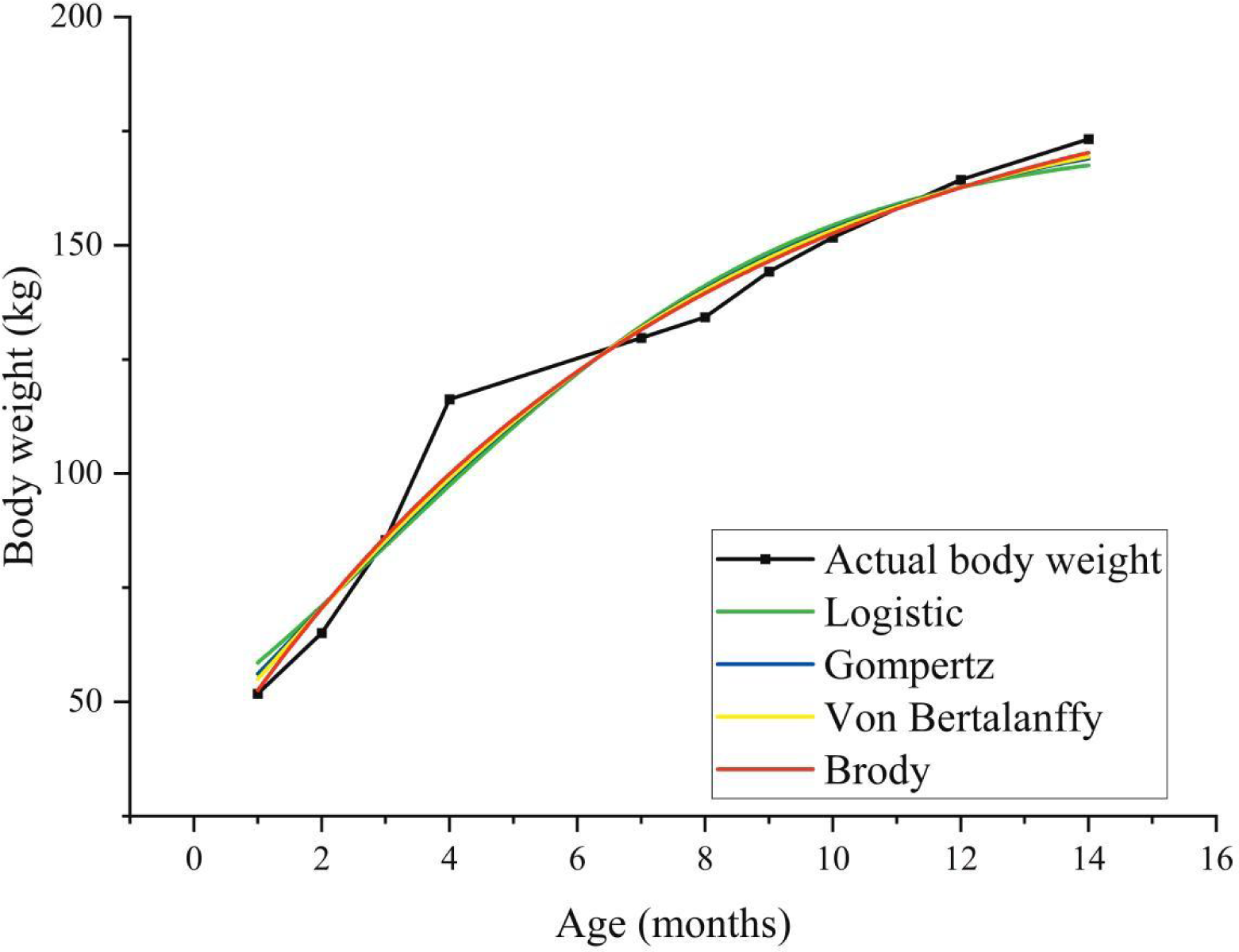
Growth curve models of body weight for female Jiangyue donkeys

**Table 11.** Body weight data of female Jiangyue donkeys at different months of age.

| Month<br>of age | Sample size | Data type | Body weight (kg) |
| --- | --- | --- | --- |
| 1 | 17 | Mean | 51.79 |
|  |  | Standard deviation | 10.74 |
| 2 | 42 | Mean | 65.07 |
|  |  | Standard deviation | 11.67 |
| 3 | 33 | Mean | 85.45 |
|  |  | Standard deviation | 11.58 |
| 4 | 28 | Mean | 116.27 |
|  |  | Standard deviation | 9.82 |
| 7 | 10 | Mean | 129.70 |
|  |  | Standard deviation | 16.20 |
| 8 | 22 | Mean | 134.23 |
|  |  | Standard deviation | 16.52 |
| 9 | 29 | Mean | 144.21 |
|  |  | Standard deviation | 16.16 |
| 10 | 29 | Mean | 151.74 |
|  |  | Standard deviation | 13.27 |
| 12 | 14 | Mean | 164.36 |
|  |  | Standard deviation | 20.17 |
| 14 | 17 | Mean | 173.26 |
|  |  | Standard deviation | 21.39 |

<sup>1)</sup> Data for 5, 6, 11 and 13 months of age were not included in the statistical analysis due to insufficient valid sample size.

**Table 12.** Fitting equation parameters and coefficient of determination of different models.

| Fitting model | Fitting parameter |  |  | R<sup>2</sup> | P value |
| --- | --- | --- | --- | --- | --- |
|  | A | B | k |  |  |
| Logistic | 173.611 | 2.668 | 0.307 | 0.96418 | <0.0001 |
| Gompertz | 180.961 | 1.457 | 0.219 | 0.99938 | <0.0001 |
| Brody | 195.057 | 0.836 | 0.135 | 0.99962 | <0.0001 |
| Von Bertalanffy | 184.410 | 0.401 | 0.190 | 0.99992 | <0.0001 |

## DISCUSSION

### Descriptive Statistical Analysis of Body Weight and Body Measurements

For donkeys, body measurements and body weight are intuitive indicators to evaluate their growth and development and production potential [12]. The coefficients of variation of body weight of growing and adult female Jiangyue donkeys were 19.04% and 16.00%, respectively, the highest among all traits. Similar results were also found in Dezhou donkeys [13] and Xinjiang donkeys [14], indicating that the body weight of donkeys had significant individual differences, and its range of extreme values was the most prominent among all traits, which could be used as an important selection index in breeding work. In addition, the rump width of growing donkeys had the largest CV (9.22%), which might be related to the unbalanced development of the hindquarters during the growing period. The chest width of adult donkeys had the largest CV (9.16%), which may reflect greater phenotypic variability in this population or indicate that this trait was greatly influenced by later feeding and management levels, thereby providing space for breeding improvement. The other body measurement traits showed stability with CVs all below 9%.

### Correlation Analysis of Body Weight and Body Measurements

Correlation analysis is used to study the correlation between variables, test their independence and positive/negative correlation, and the correlation coefficient measures the strength of their association [15]. In both the growing and adult stages, the correlation coefficient between chest circumference and body weight of female Jiangyue donkeys was the largest. Similar conclusions were obtained in the studies on Xinjiang donkeys [14], Dezhou donkeys [16], West African donkeys [17] and Miranda donkeys [18], which were consistent with the findings of this study. However, the results were not completely consistent with the study on Jiangyue donkeys by Xiao Haixia et al. [19], which might be caused by differences in the number, gender, month of age, feeding environment and other factors of the experimental donkeys. The withers height and rump height of female Jiangyue donkeys had the strongest correlation in both stages, indicating a coordinated growth pattern among height-related body dimensions during development. In contrast, the correlation of head length, neck length, cannon circumference and other traits with body weight was relatively weak in both stages, indicating that their direct contribution to body weight was lower than that of other body measurement indicators, which might be related to local skeletal structure rather than overall muscle development.

### Stepwise Regression Analysis of Body Weight and Body Measurements

Stepwise regression screens and finally retains all independent variables that have a significant impact on the dependent variable, thus constructing the optimal regression model [20]. In this study, the optimal prediction models for the body weight of growing and adult female Jiangyue donkeys were established respectively through stepwise regression analysis. The model for growing donkeys included six variables: body length, chest circumference, chest width, chest depth, rump length and rump width, with an R² of 0.918. The model for adult donkeys included three variables: body length, chest circumference and rump width, with an R² of 0.844. Body length, chest circumference and rump width were the common indicators for predicting the body weight of female Jiangyue donkeys in both growing and adult stages, so these indicators should be taken into key consideration in the selection and breeding of Jiangyue donkeys. Xiao Guoliang et al. [14] conducted a stepwise regression analysis on the body weight and body measurements of adult female Xinjiang donkeys, and the main indicators finally included in the regression equation were chest circumference and body length, with which the optimal regression model was constructed, which was consistent with the results of this study to a certain extent. John et al. [21] found in the study on donkeys in northwest Nigeria that withers height and chest circumference were the best indicators for predicting the body weight of adult female donkeys, and the optimal regression model was constructed accordingly, which was somewhat different from the results of the adult donkey model in this study. Xiao Haixia et al. [22] established the optimal regression model between body weight and body measurement traits of Jiangyue donkeys of different ages, and the finally selected indicators were withers height, body length and chest circumference. The results of this study were not completely the same, which might be related to the gender composition of the sample and the range of body measurement traits analyzed. All the above studies confirmed that body measurements had a certain predictive ability for body weight, but the indicators were different. However, it should be noted that the regression models established in this study are population-specific and require external validation with independent datasets before being generalized to other populations. One limitation of this study is that age was not included as a covariate in the regression analysis. Although we divided the study population into growth stages (growing period and adult period), thereby reducing age variation, the age span within each stage may still exert some influence on the results. Therefore, in breeding practice, scientific breeding strategies should be formulated according to the specific population and trait range to realize the continuous optimization of breeding efficiency.

### Growth Curve Fitting with Different Models

Growth curve models have been widely used to fit body weight in cattle [23], sheep [24], horses [25], poultry [26,27,28] and other animals, but the research on donkeys is relatively limited. These four nonlinear models were selected because they are among the most widely applied and well-established functions for describing sigmoidal growth patterns in livestock. The results showed that the Von Bertalanffy model had the highest goodness of fit (R²=0.99992), indicating that this model could well describe the growth and development law of female Jiangyue donkeys. Zhang et al. [29] used different models to fit the growth curve of Dezhou donkeys, and the results showed that the Brody model was the best model for fitting the growth curve of Dezhou donkeys aged 0 to 18 months. Li Xuanyue et al. [30] used the Logistic and Gompertz models to fit the growth curve of Jiangyue donkeys aged 0 to 24 months and found that the Gompertz model had the best fitting effect on the body weight growth curve. This study also found that the Gompertz model was better than the Logistic model in fitting, which was consistent with this conclusion. When the Brody and Von Bertalanffy models were further introduced for comprehensive comparison, the Von Bertalanffy model showed the best fitting effect, which indicated that the Von Bertalanffy model might be more suitable for the gradual and moderate growth characteristics of female Jiangyue donkeys aged 1 to 14 months in describing their growth curve. In practical research, the optimal fitting model should be determined through multi-model comparison. However, it should be noted that the growth curve established in this study is based on data limited to 1 to 14 months of age, and caution should be exercised against over-extrapolation when applying the model to other age stages. Although the Von Bertalanffy model showed the highest R² within the observed age range, model selection should ideally also consider additional criteria such as residual behavior, prediction error, and biological plausibility of the estimated parameters.

## CONCLUSION

This study analyzed body weight and body measurement data from 484 female Jiangyue donkeys at different growth stages. Optimal regression equations for body weight were established for growing (R²=0.918) and adult donkeys (R²=0.844) (both p<0.01). Growth curve fitting on 241 donkeys of different months of age identified the Von Bertalanffy model as the best fit (R²=0.99992). In conclusion, this study clarified the quantitative relationship between body measurements and body weight and described age-related growth patterns, providing a theoretical basis for scientific evaluation, systematic breeding, and efficient utilization of Jiangyue donkey germplasm resources.

## AUTHORS’ CONTRIBUTION

Conceptualization: Ren W, Liu L.

Data curation: Ren W, Zhao L, Yin H.

Formal analysis: Ren W.

Methodology: Ren W.

Software: Ren W, Zhao L, Yin H.

Validation: Ren W, Liu L.

Investigation: Ren W.

Writing - original draft: Ren W.

Writing - review & editing: Ren W, Zhao L, Yin H, Liu L.

## ACKNOWLEDGEMENTS

We thank all authors for their help in this research.

## SUPPLEMENTARY MATERIAL

Not applicable.

## DATA AVAILABILITY

The datasets analyzed during the current study are available from the corresponding author upon reasonable request.

## ETHICS APPROVAL

Animal experimental procedure in this study was approved by the Ethics of Xinjiang Agriculture University (Approval number: 2023040).

## DECLARATION OF GENERATIVE AI

No AI tools were used in this article.

